# A microbial metabolite, Lithocholic acid, suppresses IFN-γ and AhR expression by human cord blood CD4 T cells

**DOI:** 10.1101/491241

**Authors:** Anya Nikolai, Makio Iwashima

## Abstract

Vitamin D is a well-known micronutrient that modulates immune responses by epigenetic and transcriptional regulation of target genes, such as inflammatory cytokines. Our group recently demonstrated that the most active form of vitamin D, calcitriol, reduces expression of a transcription factor known as the aryl hydrocarbon receptor (AhR) and inhibits differentiation of a pro-inflammatory T cell subset, Th9. Lithocholic acid (LCA), a secondary bile acid produced by commensal bacteria, is known to bind to and activate the vitamin D receptor (VDR) in a manner comparable to calcitriol. In this study, Naïve CD4 T cells were isolated from healthy human umbilical mononuclear cells and were stimulated in the presence or absence of LCA. We determined the effect of LCA on human cord blood T cell activation by measuring IFN-γ production and protein expression of IFN-γ receptor-mediated signaling molecules. We found that LCA reduces production of IFN-γ and decreases phosphorylation of STAT1 as well as expression of AhR, STAT1, and IRF1 by activated human cord blood CD4 T cells. LCA inhibits expression of IFN-γ and reduces its receptor signaling molecules.

**Highlights:**

- Lithocholic acid suppresses IFNγ production by CD4 T cells.
- Lithocholic acid suppresses STAT1 and IRF1 expression by activated CD4 T cells.
- Lithocholic acid suppresses AhR in a comparable manner to calcitriol.

## 1.1 Introduction

Vitamin D deficiencies are correlated with multiple diseases, such as autoimmune disorders, hypocalcemia, and inflammatory disorders [1]. Activation of the vitamin D receptor (VDR) leads to up-or down-regulation of more than two-hundred genes [2]. Our recent study demonstrated that calcitriol, the most active vitamin D derivative, suppresses the expression of the aryl hydrocarbon receptor (AhR) by activated T cells and inhibits Th9 differentiation [3]. AhR plays pivotal roles in differentiating multiple T cell subsets and producing cytokines such as TNF-α and IFN-γ [4]. Together, these data illustrate AhR as a significant target for vitamin D to modulate the immune system.

Lithocholic acid (LCA) is a secondary bile acid metabolized by intestinal bacteria and activates VDR in a manner comparable to calcitriol [5–7]. The crystal structure of the LCA/VDR complex demonstrated that LCA and calcitriol bind the same ligand binding region of the VDR [8]. LCA is generated via 7α-dehydroxylation of the primary bile acid chenodeoxycholic acid (CDCA). To produce LCA, bacteria must express genes under the *bai* (bile acid inducible) operon. Among the bacteria in the human gut microbiome, Eubacterium and Clostridium genera express the *bai* operon [9]. Based on these data, we hypothesize that LCA functions as a bacteria-derived immunomodulatory factor that inhibits T cell activation in a manner similar to calcitriol in humans. We were particularly interested in the effect of LCA on perinatal T cells since commensal bacteria swiftly colonize newborn intestinal tracts and establish homeostasis with the host immune system [10]. Here we report the effect of LCA on activation of T cells from human umbilical cord blood.

## 1.2 Materials and Methods

### 1.2.1 Cell Culture

Naïve CD4 T cells were harvested from whole cord blood. Blood collection was performed with IRB approval (Loyola University Chicago (IRB# 203678081012)). Umbilical cord blood was collected at Loyola Gottlieb Memorial Hospital from de-identified donors that met our collection criteria (exclusion criteria: evidence of active malignancies, use of medications that affect the immune system (such as glucocorticoids and immunosuppressants), uncontrolled hyper-or hypothyroidism, presence of an autoimmune disease, and/or presence of an active infection).

Naïve CD4 T cells were obtained via negative selection using an EasySep™ Human Naïve CD4+ T Cell Enrichment Kit (Stem Cell Technologies, Vancouver, BC, Canada). All samples maintained at or above a 90% purity rating (not shown). Cells were treated once at the time of stimulation with calcitriol (10 nM; Sigma-Aldrich, St. Louis, MO) or lithocholic acid (10μM; Sigma-Aldrich), and plated on non-treated CytoOne 48 or 96 well plates (USA Scientific, Ocala, FL) that were coated with anti-CD3 (OKT3; 5μg/ml; BioLegend, San Diego, CA) and anti-CD28 (28.2; 5μg/ml; BioLegend) for cell stimulation. Control cells were treated with dimethyl sulfoxide (DMSO) since calcitriol and LCA were reconstituted in DMSO.

### 1.2.2 Western blot

Equal numbers of cells (1.0×10^6^ cells/100μl) were lysed in SDS sample buffer (2% SDS, 125mM DTT, 10% glycerol, 62.5mM Tris-HCl (pH 6.8)) and proteins were subjected to Western blot analysis using the following antibodies: anti-AhR (Santa Cruz Biotechnology, Santa Cruz, CA), anti-STAT1 (Cell Signaling Technology (CST), Danvers, MA), anti-pSTAT1 (CST), anti-IRF1 (CST), and anti-β-actin antibodies (Sigma-Aldrich). Signals were detected with the ECL system (GE Healthcare, Piscataway, NJ). The relative intensity of each band was determined by ImageJ software (National Institutes of Health) after normalization using β-actin as the control.

### 1.2.3 Cytokine expression

Cell supernatants were harvested five days post-treatment. Supernatants were analyzed for expression of T cell cytokines using the LEGENDplex Human Th cytokine panel (BioLegend) according to manufacturer’s instructions on a BD FACS CANTO II Flow cytometer (BD Biosciences, San Jose, CA). The minimum detectable concentration of this assay for both IFN-γ and TNF-α is 1.0 pg/ml. Intra-assay CVs for IFN-γ and TNF-α are 10% and 9%, respectively. Inter-assay CVs for IFN-γ and TNF-α are 12% and 8%, respectively.

### 1.2.4 Statistics

Statistical analysis was performed using GraphPad Prism software (GraphPad Software, CA). One-way ANOVA with Tukey’s posttest for multiple comparisons was used to compare cytokine concentrations.

## 1.3 Results

### 1.3.1 The effects of LCA on cytokine expression

Previous reports demonstrated that calcitriol reduces the ability of adult human T cells to produce the inflammatory cytokine IFN-γ [11]. Thus, since LCA binds and activates the VDR, we hypothesized that perinatal T cells treated with LCA would also yield a reduction in the production of IFN-γ. To test this, we stimulated naïve CD4 T cells from human cord blood in the presence or absence of either calcitriol or LCA for five days. We harvested the culture supernatant and assessed the production of cytokines by a multiplex cytokine assay system (Fig. 1a). As predicted, treatment of T cells with calcitriol or LCA significantly reduced IFN-γ expression, but we did not see a significant change in TNF-α, another Th1 like cytokine.

**Figure 1:**
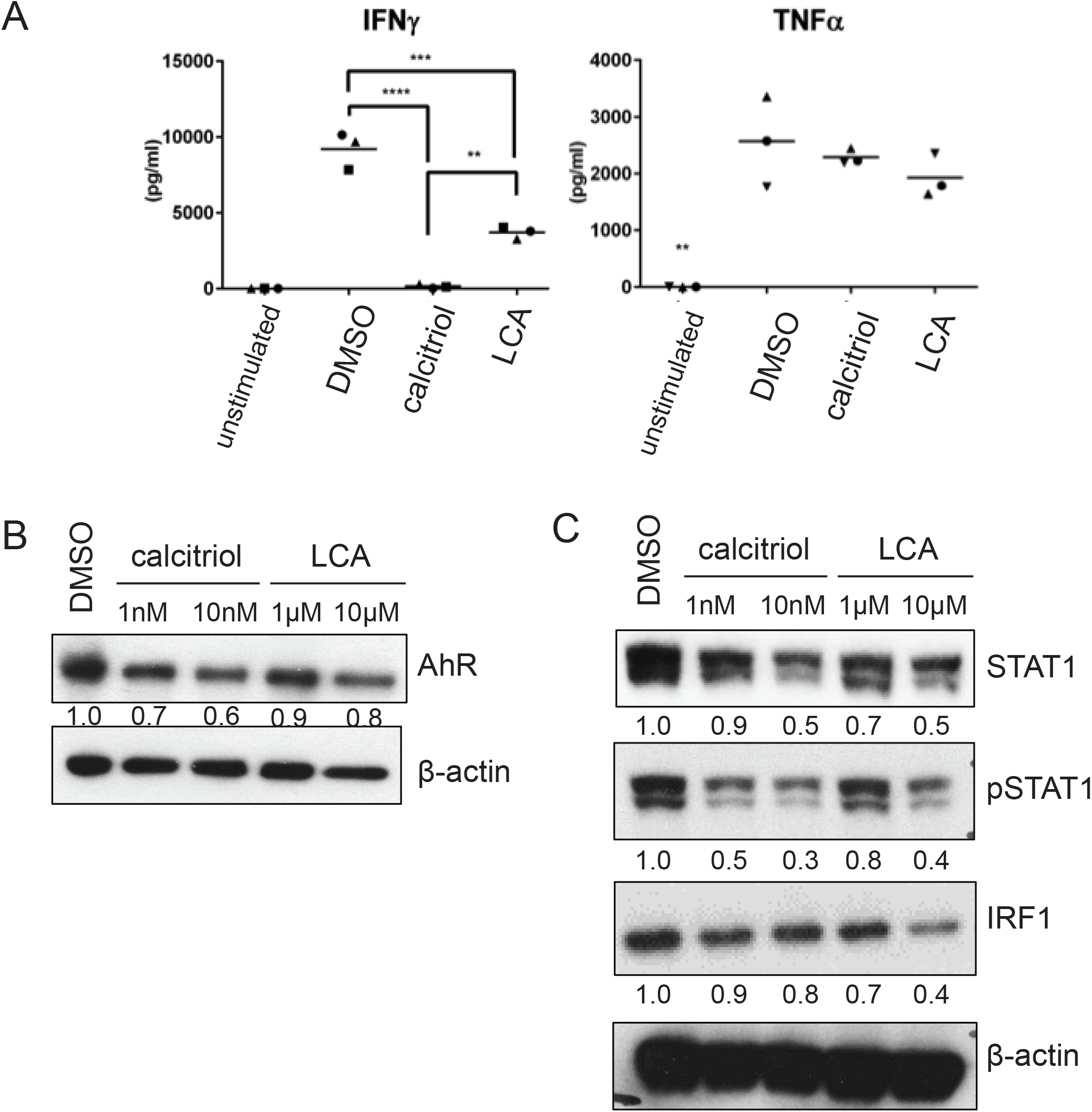
Effect of LCA on human umbilical cord blood CD4 T cells. Effect of LCA and calcitriol on (a) IFN-γ production; (b) AhR expression; and (c) STAT1, pSTAT1, and IRF1 expression. Human CD4 T cells from cord blood were stimulated by plate-bound anti-CD3/anti-CD28 antibodies and (a) supernatants were harvested five days post-treatment or (b and c) lysates were harvested 3 days post treatment. Stimulation was performed with treatment of DMSO, 10nM calcitriol, or 10μM LCA; 1nM calcitriol and 1uM LCA treated samples were also collected for lysates. Each cytokine level was determined by LEGENDplex cytokine bead array. Each symbol represents a different donor, N=3. Statistical significance was calculated using one-way ANOVA (\*\**p*<0.01; \*\*\**p*<0.001; \*\*\*\**p*<0.0001). Protein expression for (b and c) was semi-quantitatively determined by Image-J software analysis of western blot membranes. Western blot data is representative of two donor samples.

### 1.3.2 LCA mediated reduction of AhR

Our previous work showed that calcitriol reduces expression of the aryl hydrocarbon receptor (AhR) [3]. AhR plays a pivotal role in the differentiation of multiple CD4 T cell subsets including Th17, Th9, and Tregs [12]. If LCA treatment functions in a manner similar to calcitriol, then we predict that LCA also reduces protein expression of AhR. When we cultured naïve CD4 T cells in the presence or absence of LCA for 3 days, LCA reduced AhR expression in a dose-dependent manner at a level comparable to calcitriol, as expected (Fig 1b).

The data above demonstrated that LCA is a potent inhibitor of IFN-γ and AhR expression by naïve CD4 T cells. When activated, the IFN-γ receptor induces phosphorylation of STAT1. Phosphorylated STAT1 (phospho-STAT1) forms either as a homodimer, acts as a gamma-activating factor (GAF), or heterodimerizes with STAT2 [13]. Dimerized STAT1 translocates to the nucleus and upregulates transcription of IFN-γ stimulated genes such as *irf1* (interferon regulatory factor 1).

### 1.3.3 The impact of LCA on IFN-γ receptor-mediated signaling

Since LCA reduced IFN-γ production by naïve CD4 T cells, we hypothesized that the STAT1-IRF1 signaling pathway is also inhibited by LCA. As anticipated, we observed a substantial reduction in phosphorylation of STAT1 and expression of IRF-1 in T cells stimulated in the presence of calcitriol or LCA in a dose-dependent manner (Fig. 1c). We also observed a reduction in STAT1 expression. IFN-γ mediates a positive feedback loop in T cells by inducing expression of the signaling molecules that function downstream of the IFN-γ receptor, including STAT1. Thus, LCA may be able to directly suppress STAT1 and IRF-1 or this may be an indirect effect through the decrease of IFN-γ.

## 1.4 Discussion

### 1.4.1 LCA potentially reduces Th1 phenotype, may play an immunomodulatory role in the gut

Accumulating evidence suggests that there is a causative link between the gut microbiome and the intestinal innate and adaptive immune systems [14–16]. One of the key functions of the microbiome in immune homeostasis is to produce factors, such as short-chain fatty acids (SCFAs), that can modulate immune responses [17]. SCFAs stimulate cells to secrete antimicrobial peptides, reinforce the intestinal barrier, and stimulate the innate immunity to skew T cell responses. SCFAs also promote the expansion of regulatory T cells [18–20]. Our data shows that LCA is another factor produced by commensal bacteria that is immunomodulatory by inhibiting production of IFN-γ and its receptor signaling. Furthermore, IRF-1 is known to upregulate the expression of the IL-12 receptor, which can promote Th1 differentiation [21]. Our results demonstrate that LCA might inhibit T cell differentiation toward Th1 by reducing IRF-1. This positively correlates to our data that shows the Th1 cytokine, IFN-γ, is reduced. In addition, butyrate, an immunoregulatory SCFA, induces expression of the VDR [22]. An increase in VDR expression could enhance the reactivity of T cells to vitamin D as well as to LCA. Together, SCFAs and LCA could cooperatively suppress immune responses in the human intestinal environment by changing the development of effector and regulatory T cells.

### 1.4.2 Further need for studying LCA signaling in humans

This study was completed entirely *in vitro* using human umbilical cord blood. Our lab has found that multiple T cell responses are inconsistent between human and mouse T cells (not shown). Due to this, we found it more physiologically relevant to present this data using human donors, rather than a mouse model. It should be noted that LCA has a toxic effect on human tissues [14]. However, it has been reported that activation of VDR by LCA leads to the production of cytochrome 450 enzymes and subsequent detoxification of LCA [23]. This could be a natural protective mechanism against the toxicity of LCA. Due to the fact that LCA can interact with the VDR, and our findings above that LCA reduces the Th1 phenotype, LCA may reduce gut inflammation, but the concentration must stay within healthy physiological levels or it could be detrimental to the host. These dichotomies of the pro- and anti-inflammatory effects of LCA calls for further investigation of the impact of LCA on the immune system.

## 1.5 Acknowledgements

Authors thank Christina Cunha for critically reading the manuscript and the staff of Loyola Gottlieb Memorial Hospital delivery room for collecting cord blood.

This work was supported by National Institute of Health [R01AI100129] and by the Van Kampen Cardiopulmonary Research Fund.

## References

1 Yin K, Agrawal DK: Vitamin D and inflammatory diseases. J Inflamm Res 2014;7:69–87.

2 Ramagopalan SV, Heger A, Berlanga AJ, Maugeri NJ, Lincoln MR, Burrell A, Handunnetthi L, Handel AE, Disanto G, Orton SM, Watson CT, Morahan JM, Giovannoni G, Ponting CP, Ebers GC, Knight JC: A ChIP-seq defined genome-wide map of vitamin D receptor binding: associations with disease and evolution. Genome research 2010;20:1352–1360.

3 Takami M, Fujimaki K, Nishimura MI, Iwashima M: Cutting Edge: AhR Is a Molecular Target of Calcitriol in Human T Cells. Journal of immunology 2015;195:2520–2523.

4 Barrett B, Brown R, Rakel D, Mundt M, Bone K, Barlow S, Ewers T: Echinacea for treating the common cold: a randomized trial. Ann Intern Med 2010;153:769–777.

5 Begley M, Gahan CG, Hill C: The interaction between bacteria and bile. FEMS Microbiol Rev 2005;29:625–651.

6 Wells JE, Hylemon PB: Identification and characterization of a bile acid 7alpha-dehydroxylation operon in Clostridium sp. strain TO-931, a highly active 7alpha-dehydroxylating strain isolated from human feces. Appl Environ Microbiol 2000;66:1107–1113.

7 Makishima M, Lu TT, Xie W, Whitfield GK, Domoto H, Evans RM, Haussler MR, Mangelsdorf DJ: Vitamin D receptor as an intestinal bile acid sensor. Science 2002;296:1313–1316.

8 Ikura T, Ito N: Crystal Structure of the Vitamin D Receptor Ligand-Binding Domain with Lithocholic Acids. Vitam Horm 2016;100:117–136.

9 Bhowmik S, Jones DH, Chiu HP, Park IH, Chiu HJ, Axelrod HL, Farr CL, Tien HJ, Agarwalla S, Lesley SA: Structural and functional characterization of BaiA, an enzyme involved in secondary bile acid synthesis in human gut microbe. Proteins 2014;82:216–229.

10 Gensollen T, Iyer SS, Kasper DL, Blumberg RS: How colonization by microbiota in early life shapes the immune system. Science 2016;352:539–544.

11 Di Rosa M, Malaguarnera M, Nicoletti F, Malaguarnera L: Vitamin D3: a helpful immuno-modulator. Immunology 2011;134:123–139.

12 Gargaro M, Pirro M, Romani R, Zelante T, Fallarino F: Aryl Hydrocarbon Receptor-Dependent Pathways in Immune Regulation. Am J Transplant 2016;16:2270–2276.

13 Majoros A, Platanitis E, Kernbauer-Holzl E, Rosebrock F, Muller M, Decker T: Canonical and Non-Canonical Aspects of JAK-STAT Signaling: Lessons from Interferons for Cytokine Responses. Frontiers in immunology 2017;8:29.

14 Sun J, Chang EB: Exploring gut microbes in human health and disease: Pushing the envelope. Genes Dis 2014;1:132–139.

15 Purchiaroni F, Tortora A, Gabrielli M, Bertucci F, Gigante G, Ianiro G, Ojetti V, Scarpellini E, Gasbarrini A: The role of intestinal microbiota and the immune system. Eur Rev Med Pharmacol Sci 2013;17:323–333.

16 Woo V, Alenghat T: Host-microbiota interactions: epigenomic regulation. Curr Opin Immunol 2017;44:52–60.

17 Geuking MB, McCoy KD, Macpherson AJ: Metabolites from intestinal microbes shape Treg. Cell Res 2013;23:1339–1340.

18 Furusawa Y, Obata Y, Fukuda S, Endo TA, Nakato G, Takahashi D, Nakanishi Y, Uetake C, Kato K, Kato T, Takahashi M, Fukuda NN, Murakami S, Miyauchi E, Hino S, Atarashi K, Onawa S, Fujimura Y, Lockett T, Clarke JM, Topping DL, Tomita M, Hori S, Ohara O, Morita T, Koseki H, Kikuchi J, Honda K, Hase K, Ohno H: Commensal microbe-derived butyrate induces the differentiation of colonic regulatory T cells. Nature 2013;504:446–450.

19 Arpaia N, Campbell C, Fan X, Dikiy S, van der Veeken J, deRoos P, Liu H, Cross JR, Pfeffer K, Coffer PJ, Rudensky AY: Metabolites produced by commensal bacteria promote peripheral regulatory T-cell generation. Nature 2013;504:451–455.

20 Singh N, Gurav A, Sivaprakasam S, Brady E, Padia R, Shi H, Thangaraju M, Prasad PD, Manicassamy S, Munn DH, Lee JR, Offermanns S, Ganapathy V: Activation of Gpr109a, receptor for niacin and the commensal metabolite butyrate, suppresses colonic inflammation and carcinogenesis. Immunity 2014;40:128–139.

21 Kimura A, Naka T, Nakahama T, Chinen I, Masuda K, Nohara K, Fujii-Kuriyama Y, Kishimoto T: Aryl hydrocarbon receptor in combination with Stat1 regulates LPS-induced inflammatory responses. The Journal of experimental medicine 2009;206:2027–2035.

22 Daniel C, Schroder O, Zahn N, Gaschott T, Steinhilber D, Stein JM: The TGFbeta/Smad 3-signaling pathway is involved in butyrate-mediated vitamin D receptor (VDR)-expression. J Cell Biochem 2007;102:1420–1431.

23 Cheng J, Fang ZZ, Kim JH, Krausz KW, Tanaka N, Chiang JY, Gonzalez FJ: Intestinal CYP3A4 protects against lithocholic acid-induced hepatotoxicity in intestine-specific VDR-deficient mice. J Lipid Res 2014;55:455–465.

